# Parkinson’s disease medication alters rat small intestinal motility and microbiota composition

**DOI:** 10.1101/2021.07.19.452878

**Authors:** Sebastiaan Pieter van Kessel, Amber Bullock, Gertjan van Dijk, Sahar El Aidy

## Abstract

Parkinson’s disease (PD) is known to be associated with altered gastrointestinal function and microbiota composition. Altered gastrointestinal function is key in the development of small intestinal bacterial overgrowth, which is a comorbidity often observed in PD patients. Although PD medication could be an important confounder in the reported alterations, its effect *per se* on the microbiota composition or the gastrointestinal function at the site of drug absorption has not been studied. To this end, healthy (i.e., not PD model) wild-type Groningen rats were employed and treated with dopamine, pramipexole (in combination with levodopa/carbidopa), or ropinirole (in combination with levodopa/carbidopa) for 14 sequential days. Rats treated with dopamine agonists showed a significant reduction in the small intestinal motility and an increase in bacterial overgrowth in the distal small intestine. Importantly, significant alterations in microbial taxa were observed between the treated and vehicle groups, analogous to the changes previously reported in human PD vs HC microbiota studies. These microbial changes included an increase in *Lactobacillus* and *Bifidobacterium*, and decrease in *Lachnospiraceae* and *Prevotellaceae*. Importantly, certain *Lactobacillus* species correlated negatively with the plasma levels of levodopa. Overall, the results highlight the significant effect of PD medication *per se* on the gut microbiota, and the disease-associated comorbidities, including gastrointestinal dysfunction and small intestinal bacterial overgrowth.

**Research in context:** *Evidence before this study:* In the last years many studies focused on the faecal microbiota profiles of Parkinson’s disease (PD) patients and compared them to healthy control (HC) subjects and reported a difference between the microbiota profiles. Although some bacterial taxa have been reported to be differential abundant across studies there is a low consensus between the overall changes observed. It remains to be elucidated whether and how the disease status itself, the PD medication or the gastrointestinal dysfunction, an often reported comorbidity, plays a role in the altered microbiota profiles of Parkinson’s disease patients.

*Added value of this study:* Two major confounding factors potentially affecting the microbiota profiles differentiating PD patients from HC subjects are the PD medication and gastrointestinal dysfunction. We showed that PD medication elicits an decreasing effect on the small intestinal gut motility in a healthy (i.e. non Parkinson) rat model which was a contributing factor to the altered microbiota profile observed. We found that taxa reported previously to be differentially abundant were mirrored in our healthy rat model showing the impact of PD medication on microbiota profiles.

*Implications of all the available evidence:* This study showed that PD medication and gastrointestinal motility are important factors in the microbiota profiles and could explain some of the differential abundant taxa reported in the cross-sectional PD microbiota studies. Future studies should take the potential side effects of medication that could elicit alterations of the microbiota composition into account when seeking for microbial biomarkers of disease status.

## Introduction

The microbiota composition of patients with Parkinson’s disease (PD) have been investigated in various, mainly cross-sectional, studies over the past years, in comparison to matching-healthy control (HC) subjects.^1–16^ However, there is low consensus among the findings in these reports. Recently, the sequencing data from most of these studies have been pooled and re-analysed in order to identify common alterations in the gut microbiota of PD patients.^17^ Romano *et al*. concluded that it was impossible to determine whether the changes in microbiota composition were causally linked to the disease *per se* in the pooled data analysis due to confounding factors such as PD medication, which was considerably variable among the studies included.^17^ Indeed PD medication is a major differentiating factor between PD patients and HC subjects and only a few studies^1,6,9,12,14^ investigated or reported the effect of medication on the microbiota profiles in PD patients. So far, no studies were performed in healthy subjects, which is required to determine whether the PD medication *per se* affects the microbiota composition.

Most PD medication work through their effect on the dopaminergic system in the brain. The medication, in turn, intrinsically influence the peripheral dopaminergic systems in the enteric nervous system (ENS) ^18,19^, and the immune system.^20,21^ Dopamine and/or dopamine agonists are known to affect the gut motility in rodents, dogs, and humans^22^ (and references in there^23–36^). Gut motility is usually inferred by Bristol-Stool Score, which is known to be a major contributor to the variation in the fecal microbiota composition.^37^ Many PD patients experience non-motor symptoms, including gastrointestinal (GI) dysfunction, such as decreased small intestinal motility^38,39^, which is one of the causes of small intestinal bacterial overgrowth (SIBO).^40^ SIBO is prevalent (up to 54·5 %) and significantly higher in PD patients.^41–43^ Recently, PD medication has been also associated with the development of GI-symptoms ^44^ or GI-transit ^45^ in PD patients. Analogously, we showed that the unabsorbed residues of levodopa that reach the distal small intestine is converted to a bioactive molecule, which reduces ileal contractility in mice *ex vivo*.^46^ Nonetheless, whether PD medication *per se* is also associated with alterations in microbiota composition, small intestinal gut motility, and SIBO remains unknown.

In this study, we show that the commonly prescribed PD medication; pramipexole and ropinirole in combination with levodopa/carbidopa, when administered to healthy rats, have a profound effect on the small intestinal motility and alterations in the microbiota composition similar to those reported in PD patients, irrespective to any PD symptoms.

## Methods

### Rat experiments

All animal procedures were approved by the Groningen University Committee of Animal experiments (approval number: AVD1050020197786) and were performed in adherence to the NIH Guide for the Care and Use of Laboratory Animals.

Thirty-six adult male WTG rats (Groningen breed, age 22 to 27 weeks) housed 2 to 4 animals/cage had ad libitum access to water and food (Altromin 1414) in a temperature (21 ± 1 °C) and humidity-controlled room (~60% relative humidity), with a 12-h light/dark cycle.

The rats were trained on drinking 10% (w/v) sucrose solution from a burette with spout as followed. On 9-13 occasions over a period of 2-3 weeks, rats were taken from their social housing cage in the beginning (within 1 hour) of the dark-phase cycle and placed in an individual training cage (L × W × H = 25 × 25 × 40 cm), without bedding, food, or water. Ten minutes after transfer to the training cages, rats were given a drinking burette with a 2·5 −mL sucrose solution (10% w/v). On 2-4 training occasions, 1·2 % carmine red (C1022, Sigma) was added to the sucrose solution. Over the course of training, all rats were trained to drink the sucrose solution avidly. After 2-3 weeks, when the training was complete, animals were designated at random to four different treatments groups, dopamine (D, n=10), pramipexole/levodopa/carbidopa (P, n=10), ropinirole/levodopa/carbidopa (R, n=10), and vehicle (VH, n=6) animals were at least from 2-3 different cages, but treated per cage because of coprophagy. Rats in the designated groups were treated for 14 consecutive days with on average 1·5 mg/kg dopamine (H8502, Sigma), 0·0625 mg/kg pramipexole (A1237, Sigma) with 7·5/1·875 mg/kg levodopa/carbidopa (D9628/C1335, Sigma), 0·15 mg/kg ropinirole (R2530, Sigma) with 7·5/1·875 mg/kg levodopa/carbidopa, or 10% sucrose (w/v) solution only (VH). Based on a person weighing 80 kg, the dosages are equivalent to 600/150 mg levodopa/carbidopa, 5 mg/day pramipexole, 12 mg/day ropinirole or 120 mg/day dopamine based on 10% from a high levodopa dose (1200 mg/day). On the last treatment day, animals were sacrificed 18·5±0·68 minutes after they had started drinking the PD medication (no differences in time of sacrifice were observed between groups: D, 18·55±0·69; P, 18·68±0·80; R, 18·4±0·67; VH, 18·28±0·47; One-way-ANOVA statistics, F=0·4977, P-value= 0·6865). All rats received their dose supplemented with 1·2% (w/v) carmine red to determine their small intestinal motility, and rats for the D and VH groups received instead of their original dose on average 7·5/1·875 mg/kg levodopa/carbidopa in order to determine the potential levodopa uptake differences between treated groups. The rats were anesthetized (by isoflurane inhalation anaesthesia) and a blood sample was taken (by heart puncture), on average 18·5±0·68 minutes after the rats had start drinking continuously from the burette (2-3 minutes of drinking). No differences in time of heart puncture were observed between groups (D, 18·55±0·69; P, 18·68±0·80; R, 18·4±0·67; VH, 18·28±0·47; One-way-ANOVA statistics, F=0·4977, P-value= 0·6865). Blood withdrawn by heart puncture was dispensed in tubes pre-coated with EDTA with a final concentration of 5 mM and stored on ice during the experiment. The collected blood samples were centrifuged at 1500× g for 10 min at 4 °C and the plasma was stored at −80 °C prior to catecholamine extraction. Then rats were killed by decapitation (using a rodent guillotine), and the small intestine from stomach to cecum was taken from abdominal cavity and dissected, where the first 5 cm was considered as duodenum, the remaining part (jejunum and ileum) were dissected in 6 equal pieces and their luminal contents were collected from every section by gentle pressing and were stored on ice during the experiment. Directly after, the samples were used for carmine red determination and colony forming unit (CFU) counting, as described below. After the samples were processed they were snap frozen in liquid N_2_ and stored at −80 °C.

### Carmine red assay

Part of the luminal content per small intestinal section was suspended in DMSO 20% (w/v) were vortexed vigorously and 80μl was distributed in a 96 well plate. Spectrum from 450-800 (10 nm/step) was measured (carmine red has 2 peaks at 530/570). Because of high background differences, the spectrum was linearized between 510 and 590 nm using a fitted line (y= a * x + b). The slope (a) and the intercept (b) were calculated using the data points from 510 and 590 nm, and the calculated value (x) for 570 nm (y) was subtracted from the measured value. Next, because the animals were not fasting before the treatment, the linearized values were scored binary, a score of 1 was given when the value was larger than the threshold of 0·003. Finally the geometric centre, concluded to be the most sensitive and reliable measure of intestinal transit^47^, was calculated by multiplying the binary score by the segment number (1 to 7, from end of stomach to beginning cecum).

### Colony Forming Unit assay

Contents from the jejunal segments and ileal segments were mixed and suspended in GM17/17% glycerol media to preserve the bacterial viability after storing at −80 °C. The suspended jejunal and ileal contents were 10-fold serial diluted in PBS and 10 μl was spotted in triplicates on chopped meat media plates (CMM; beef extract, 10 g/L; casitone, 30 g/L; yeast extract, 5 g/L; K_2_HPO_4_, 5 g/L, menadione, 1 μg/mL, cysteine, 0·5 g/L; hemin 5 μg/mL, 15 g/L agar), which were incubated for 48 hours aerobically and anaerobically (1·5% H_2_, 5% CO_2_, balance with N_2_) in a Coy Laboratory Anaerobic Chamber (neo-Lab Migge GmbH, Heidelberg, Germany) at 37 °C before colony forming units were counted.

### Catecholamine extraction

Plasma samples were thawed on ice and a spatula-tip (~5mg) of activated alumina powder (199966, Sigma) was added to each well of a 96-well AcroPrep filter plate with 0·2 μM wwPTFE membrane (514-1096, VWR). A100 μL of plasma sample, 1 μM DHBA (3,4-dihydroxybenzylamine hydrobromide, 858781, Sigma) as an internal standard, and 800 μL of TE buffer (2·5% EDTA; 1·5 M Tris/HCl pH 8·6) were added sequentially to the wells. Liquid was removed using a 96-well plate vacuum manifold and the alumina were washed twice with 800 μl of H_2_O. Catechols were eluted using 0·7% HClO_4_, which was incubated for 30 min at RT. Samples were injected in a HPLC-ED system (Ultimate 3000 SD HPLC system coupled to Ultimate 3000 ECD-3000RS electrochemical detector with a glassy carbon working electrode (DC amperometry at 800 mV), Thermo Scientific). Samples were analysed on a C18 column (Kinetex 5 μM, C18 100 Å, 250 × 4·6 mm, Phenomenex, Utrecht, The Netherlands) using a gradient of water/methanol with 0·1% formic acid (0–3 min, 99% H_2_O; 3–7 min, 99–30% H_2_O; 7–10 min 30–5% H_2_O; 10–11 min 5% H_2_O; 11–18 min, 99% H_2_O). Data recording and analysis were performed using Chromeleon software (version 6·8 SR13). Potential intake differences of levodopa were corrected by using carbidopa as an internal standard.

### Levodopa decarboxylation activity test

Samples stored at −80 °C in GM17/17% glycerol were thawed ice and 300 μL of 10% (w/v) jejunal or ileal suspensions were washed once with 1 mL of ice-cold PBS to remove levodopa (given during the treatment) and glycerol from the storage medium. Pellets were resuspended in 600 μL EBB (as described before^48^) supplemented with 20μg/mL kanamycin (EBB/K) resulting in a 5% (w/v) suspension. A100 μM of levodopa was added to the suspensions and samples were incubated anaerobically (1·5% H_2_, 5% CO_2_, balance with N_2_) in a Coy Laboratory Anaerobic Chamber (neo-Lab Migge GmbH, Heidelberg, Germany) at 37 °C. Samples of 100 μl were taken at 0 and 24 h and 400 μL of methanol was added. Cells and protein precipitates were removed by centrifugation at 20,000 × g for 10 min at 4 °C. Supernatant was transferred to a new tube and the methanol fraction was evaporated in a Savant speed-vacuum dryer (SPD131, Fisher Scientific, Landsmeer, The Netherlands) at 60 °C for 90 min. The aqueous fraction was reconstituted to 0·5 mL with 0·7% HClO_4_. Samples were filtered and injected into the HPLC system described above. Dopamine and levodopa concentrations were quantified from the 24 h samples and the ratio between dopamine and levodopa was calculated to determine levodopa decarboxylation activity.

### DNA isolation and Sequencing

DNA isolation was performed based on repeated beat beating (RBB) protocol described in.^49,50^ Approximately 150-200 mg of jejunal or ileal content was weighted in screw-cap tubes containing ~0·5 g 0·1 mm glass/silica beads and 3 large 3 mm glass beads. Bacterial cells were lysed by adding 750 μL lysis buffer (NaCl, 500 mM; Tris-HCL at pH 8, 50 mM; EDTA, 50 mM; SDS, 4 % (w/v)) with sequential bead-beating 3 × 1 min with 1 min intervals on ice in a mini bead-beater (Biospec, Bartlesville, USA). Samples were incubated for 15 min, with regular mixing, at 95 °C, placed for 5 min on ice, and centrifuged at 20,000 × *g* for 30 min at 4 °C. Approximately 600 μL of the samples was recovered, centrifuged again for 5 min, before 550 μl was transferred to a new tube containing 200 μL, 10 M ammonium acetate and mixed. Samples were incubated for 5 min at ice before centrifugation at 20,000 × *g* for 30 min at 4 °C. Approximately 700 μl was transferred to a new tube, centrifuged again for 5 min before 650 μL of supernatant was transferred to a new tube containing 650 μL 2-propanol and mixed. Samples were incubated for 30 min at ice and centrifuged at 20,000 × *g* for 15 min at 4 °C. Pellets containing the DNA were washed twice with 800 and 500 μL 70% (v/v) ethanol by centrifugation at 20,000 × *g* for 10 min at 4 °C. The supernatant was discarded and the pellet was dried to air in a 37 °C heat block for 30 min. After drying, the pellets were dissolved in 200 μL TE buffer (1 mM, EDTA; 10 mM Tris-HCl at pH 8) by vortexing and incubating at 65 °C for 10 min. DNA extracted samples were stored at −80 before further clean up with the Genomic DNA Clean & Concentrator (gDCC) kit (D4011, Zymo Research, BaseClear Lab Products, The Netherlands). Samples were thawed at RT and to 0·1 mg/ml RNAse A (EN0531, Thermo Scientific) was added and incubated for 15 min at 37 °C before the clean-up with the gDCC kit. Added ChiP Binding Buffer to the RNAse A treated samples (2:1), mixed, and transferred the mixture to the gDDC column, which was centrifuged at 14,000 × *g* for 30 sec at RT. DNA bound the column was washed twice at 14,000 × *g* for 60 sec at RT with wash buffer before eluted in pre-heated (65 °C) elution buffer which was incubated for 3 min on the column. DNA integrity was checked on agarose gel before samples were outsourced for 16s (region V3-V4) amplicon metagenomic sequencing by Novogene Co., Ltd..

16S rRNA gene regions V3-V4 were amplified with primers 314F (5’-CCTAYGGGRBGCASCAG-’3) and 806R (5’-GGACTACNNGGGTATCTAAT-3’) with Phusion High-Fidelity PCR Master mix (New England Biolabs) and amplified products were verified using Agilent 5400 Fragment analyser, which all passed the quality control. PCR products were equally mixed and purified with Qiagen Gel Extraction kit before libraries for paired-end 250bp Illumina sequencing were prepared with NEBNext Ultra DNA Library Prep Kit (New England Biolabs).

### Data analysis

Paired-end reads were assigned to their samples and the barcodes and primer-sequence were truncated before merging using FLASH (V1.2·7).^51^ Quality filtering was performed as described here^52^ using QIIME (V1.7·0).^53^ Chimera sequences were removed using the UCHIME algorithm (with “Gold” database).^54^ Finally OTU calling was performed using UPARSE (v7.0·1001)^55^ and sequences with ≥97% similarity were assigned to the same OTUs. Mothur software^56^ was used for species annotation at each taxonomic rank (threshold: 0·8-1) against the SILVA Database^57^ and the phylogenetic tree was constructed using MUSCLE (Version 3.8·31).^58^

The OTU table and phylogenetic tree were imported in the R package *phyloseq* (v1·32·0).^59^ Richness and diversity were estimated on the raw OTU-counts table using *phyloseq*. For further data analysis the OTU-counts were normalized using the cumulative-sum scaling normalization (CSS) method using the R package *metagenomeSeq* (v1·30·0)^60^ and taxa were agglomerated on genus level using *phyloseq*. Unweighted UniFrac^61^ distances were calculated in *phyloseq* using the phylogenic tree rooted on the longest branch using the *root* function from R package *ape* (v5·4-1).^62^

### Statistical analyses

Data and statistical analyses were performed in GraphPad Prism (v7·0), IBM SPSS Statistics (v 26) or R (v4.0·4) in Rstudio (v1.2·5042). The One-way ANOVAs followed by Fisher’s LSD test with FDR correction in **Figure 1 and 2** were performed in GraphPad Prism. For the CFU data outliers were determined with the ROUT method (Q=0·1%) and removed using GraphPad Prism. The One-way ANOVA followed by Dunnet’s test in **Figure 4AB** was performed in SPSS. Principal Coordinate Analysis (PCoA), Principal Component Analysis (PCA) and Correspondence Analysis (CA) were performed in *phyloseq*. PERMANOVA (Permutational Multivariate Analysis of Variance) was performed using the *adonis2* function from the R package *vegan* (v2·5-6)^63^ and environmental vector fitting was performed using the function *envfit* from *vegan*. For differential abundance analysis the R package *metagenomeSeq* was used for the zero-inflated log-normal model^60^ and LDA Effect Size (LEfSe)^64^ analysis was performed in the Galaxy web application (http://huttenhower.sph.harvard.edu/galaxy/). Specific tests and significance are indicated in the figure legends.

**Figure 1.**
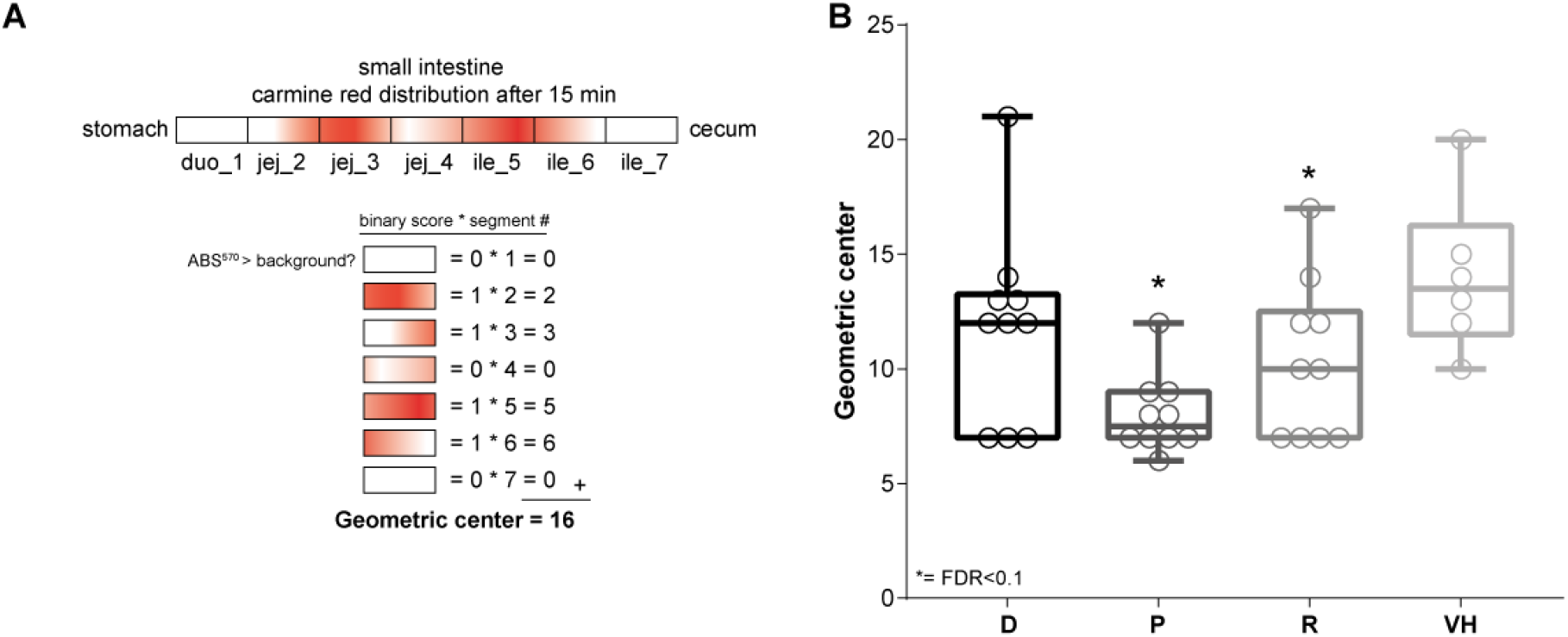
Small intestinal motility is affected by PD medication treatment. (**A**) schematic representation of the small intestine. Each rectangle represents the different sections assessed of the small intestine, where carmine red distribution in the small intestine is depicted in red. Each segment was scored binary and multiplied by the segment number resulting in the geometric centre. Duo, duodenum; jej, jejunum; ile, ileum (**B**) the geometric centre per treated group is depicted. D, dopamine; P, pramipexole; R, ropinirole; VH, vehicle (10% sucrose). Boxes represent the median with interquartile range, and whiskers represent the maxima and minima. Significance compared to VH (asterisks) was tested with One-way-ANOVA followed by Fisher’s LSD test with FDR correction.

**Figure 2.**
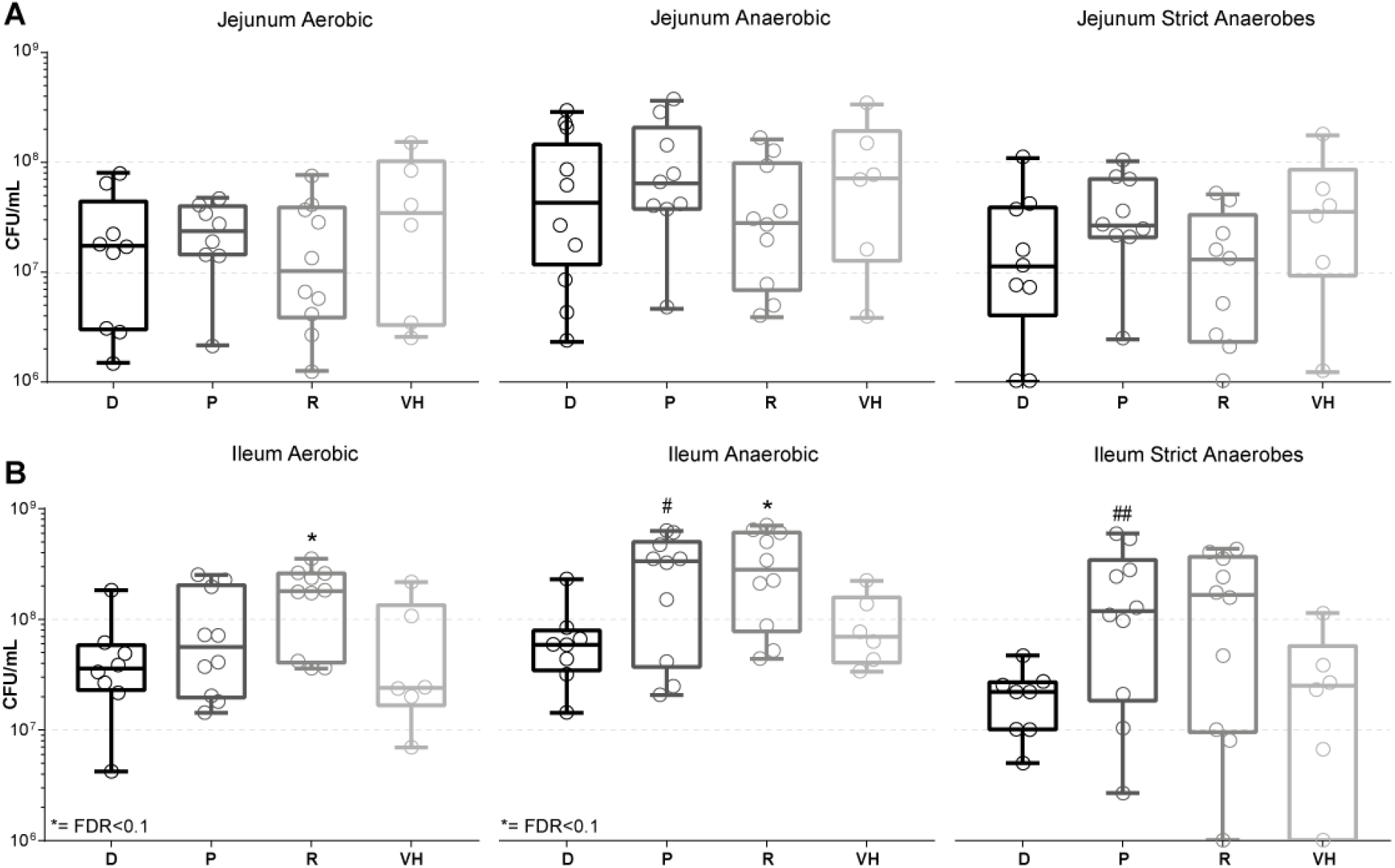
Significantly higher bacterial counts in the ileum of rats treated with PD medication. (**A**) the jejunal and (**B**) ileal colony forming units (CFUs)/ml were counted after 48 h aerobic or anaerobic incubation of jejunal and ileal content. Panels (left to right) depict the aerobic, anaerobic and strict anaerobic (anaerobic - aerobic counts) CFU/mL, respectively. D, dopamine; P, pramipexole; R, ropinirole; VH, vehicle (10% sucrose). Boxes represent the median with interquartile range, and whiskers represent the maxima and minima. Extreme outliers were removed using the ROUT method (Q=0·1%). Significance compared to VH (asterisks) was tested with One-way-ANOVA followed by Fisher’s LSD test with FDR correction (q<0·1). #, p = 0·054, q = 0·080; ##, p = 0·046, q = 0·138.

**Figure 3.**
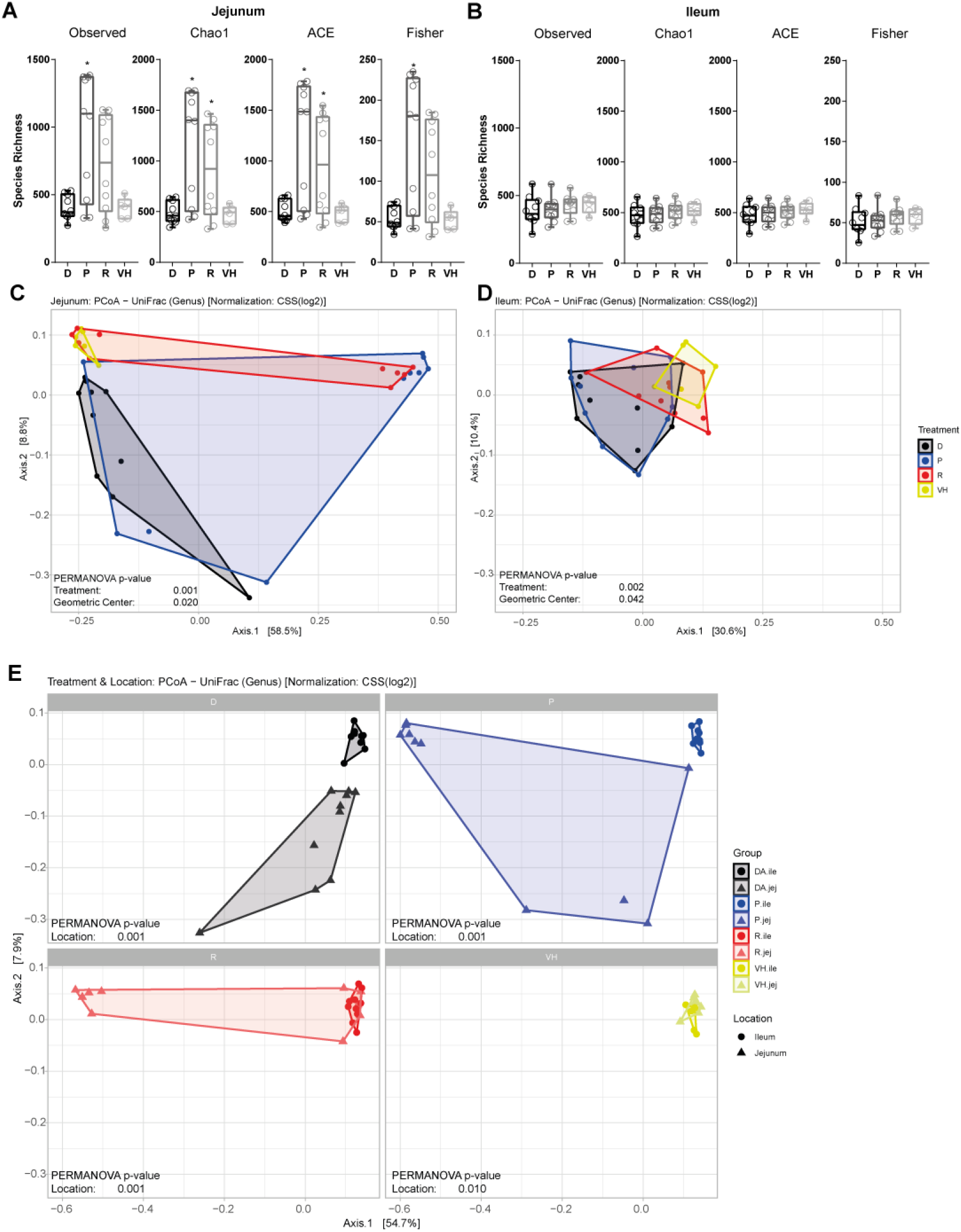
PD medication and geometric centre are significantly contributing to the variation in the microbiota composition. (**A,B**) represent the species richness (Observed, Chao1, ACE and Fisher) of the jejunum (**A**) and ileum (**B**). Boxes represent the median with interquartile range, and whiskers represent the maxima and minima. Significance compared to VH (asterisks) was tested with One-way-ANOVA followed by Fisher’s LSD test with FDR correction. (**C,D**) depict a Principal Coordinate Analysis (PCoA) using unweighted UniFrac distances at genus level using CSS scaled data of jejunum (**C**) and ileum (**D**). (**E**) depicts a PCoA using unweighted UniFrac distances at genus level using CSS scaled data of jejunum and ileum (faceted by treatment). D, dopamine; P, pramipexole; R, ropinirole; VH, vehicle (10% sucrose). Significant contribution of the variables to the variance of the PcoA was tested with Permutational Multivariate ANOVA (PERMANOVA).

**Figure 4.**
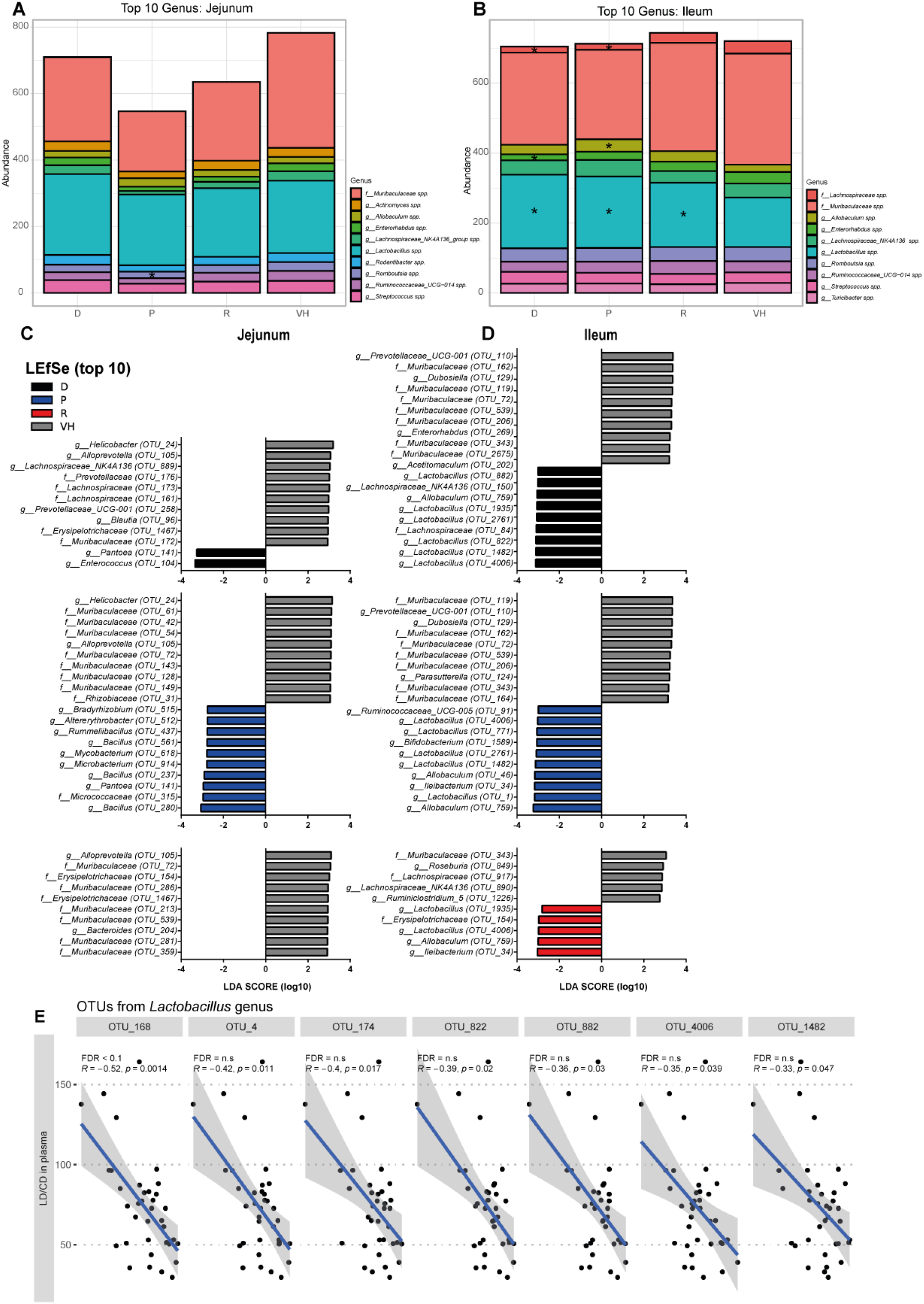
Differential abundance analysis of bacterial species among the different treatment groups and negative association with levodopa uptake. (**A,B**) represent a stacked bar plot with mean genus levels using CSS scaled data from the top 10 taxa are from jejunum (**A**) and ileum (**B**). The asterisks indicate statistical significance compared to VH group tested using one-way-ANOVA followed with Dunnett’s test. (**C,D**) represent LEfSe analysis (top 10) of the different treated groups of jejunum (**C**) and ileum (**D**). Significance was tested using one-way-ANOVA followed by a Kruskal-Wallis (KW) test and Linear Discriminant Analysis (LDA). A significant feature was considered when KW p-value < 0·01 and Log(LDA score) > 2. For all the significant extracted features see **Supplementary Table 1**. (**E**) illustrates graphs with linear models and spearman correlations of the significant OTUs of the top 50 abundant OTUs with the levodopa uptake. Only *Lactobacillus* OTU_168 remained significant after FDR correction. For the not significant OTUs see **Supplementary Table 2**. D, dopamine; P, pramipexole; R, ropinirole; VH, vehicle (10% sucrose).

## Results

### Parkinson’s disease medication affects small intestinal motility in wild-type Groningen rats

To test whether the commonly prescribed PD medication affects the small intestinal motility, wild-type Groningen (WTG) rats were employed and were treated for 14 sequential days with dopamine (D), pramipexole (P, in combination with levodopa/carbidopa as co-prescribed for PD patients), ropinirole (R, in combination with levodopa/carbidopa), or vehicle (VH). Although dopamine is not used as a treatment for PD, it was included in the study for two reasons; 1) as a control for the dopamine agonist groups, 2) PD patients usually have a higher exposure (2·5-40 fold) to dopamine, a metabolite of the levodopa treatment, than HC subjects.^65–67^ On the last treatment day, animals were sacrificed 18·5±0·68 minutes after they had started drinking the PD medication. The small intestine was sectioned into 7 pieces and their contents were assessed for carmine red spectrophotometrically. Carmine red detection was scored binary per segment (detection scored 1; no detection scored 0) and the geometric centre, a sensitive and reliable measure of intestinal transit^47^ was determined (**Figure 1A**). Pramipexole- and ropinirole-treated groups showed a significant decrease in the geometric centre compared to the vehicle, indicative of a reduced small intestinal motility (**Figure 1B**). In contrast, the dopamine-treated group did not show a significant effect on the small intestinal motility although 8/10 (up to the 3^rd^ quartile) of the points were below the median of the vehicle group. These findings suggest that PD treatment affects the small intestinal motility, which may, in turn, influence the bacterial composition and cause an increase in SIBO at the site of drug absorption.

### Increased bacterial counts in the small intestine of rats treated with Parkinson’s disease medication

Because altered small intestinal motility is known to affect the bacterial loads^40^, we determined the colony-forming units (CFU) in the jejunal and ileal content, respectively (**Figure 2AB**). The content was spotted on chopped meat media plates and were incubated aerobically and anaerobically for 48 h at 37 °C before CFUs were counted. In the jejunum, no significant differences were observed between the treated and vehicle groups (**Figure 2A**). In contrast, there was a significant increase in bacterial counts in the ileal content of the treated groups compared to the vehicle (**Figure 2B**). The ropinirole-treated animals had significantly higher bacterial counts on both the aerobic (p = 0·024, q = 0·073) and anaerobic (p = 0·020, q = 0·061) incubated plates (**Figure 2B**). Subtracting the aerobic counts from the anaerobic counts, indicative of strict anaerobic counts, showed no significant difference suggesting that the facultative anaerobes are contributing to the bacterial increase observed in the ropinirole-treated group (**Figure 2B**). In the pramipexole-treated animals, a borderline (p = 0·054, q = 0·080) increase was observed in the anaerobically incubated plates (**Figure 2B**). Similarly, the strict anaerobic counts showed a borderline increase in bacterial counts (p = 0·046, q = 0·138) (**Figure 2B**), indicating that strict anaerobes in the pramipexole-treated group may contribute to the observed bacterial higher counts. Collectively, the results imply that the reduced gut motility caused by PD medication are plausibly associated with the observed increase in bacterial counts in the ileum of the treated groups.

### Parkinson’s disease medication affects the microbiota composition, potentially through gut motility

Next, we investigated whether the PD medication resulted in changes in the small intestinal microbiota composition directly or indirectly through the altered small intestinal motility (**Figure 1B**). To this end, we performed amplicon metagenomic sequencing on the V3-V4 regions of the bacterial 16s genes. Interestingly, the richness (i.e., the number of different species observed) in the jejunum, but not in the ileum, was significantly different in the pramipexole- and ropinirole-treated groups compared to the vehicle (**Figure 3A-B**). The Shannon’s and Simpson’s diversity indices (i.e., species richness considering species proportional abundances) showed marginal changes in the jejunum between the treated and vehicle groups (**Supplementary Figure 1A-B**). In ileum, the Simpson’s index, but not inverse Simpson’s index, was significantly higher in the ropinirole-treated group compared to the vehicle (**Supplementary Figure 1B**). The data highlight an increase in the species richness only in the jejunum upon treatment with dopamine agonists, while the diversity did not change along the small intestine.

To determine whether the jejunal and ileal microbiota compositions are distinct from each other upon administration of PD medication, ß-diversity analyses using Principal Coordinate analysis (PCoA) with UniFrac distance was performed. The analysis revealed a significant difference within each treated group in both jejunal and ileal microbiota composition, respectively (**Supplementary Figure 1C-F**). The outcome showed differences in the microbial composition between the tested small intestinal locations, independent of the treatment. Since the microbiota composition is known to be different between the jejunum and ileum, we separated the analysis based on the location. The analysis showed that the treatment had a significant effect on the microbiota composition in both jejunum (PERMANOVA: R^2^= 0·223, p= 0·001) and ileum (PERMANOVA: R^2^= 0·197, p= 0·002) (**Figure 3C-D**).

Because the gut motility was significantly affected in the dopamine agonist-treated groups (**Figure 1B**), the geometric centre was also tested for its contribution in the observed changes in the microbiota composition. Indeed, the changes in geometric centre contributed significantly to the altered microbiota in the jejunum and ileum (PERMANOVA: R^2^= 0·1, p=0·020 and R^2^= 0·06, p=0·042 respectively) (**Figure 3C-D**). Additionally, correspondence analysis (CA), another ordination technique used in exploring microbial ecology, was performed and fitted with the treatment and geometric center as dependent variables (**Supplementary Figure 1G-H**). For both jejunum and ileum, treatment was a significant factor (envfit: R^2^= 0·341, p-value= <0·001 and R^2^= 0·196, p=0·019 respectively). Importantly, the changes in gut motility measured by geometric centre were strongly associated with the direction of the vehicle (envfit: p-value=0·009, R^2^= 0·263 **Supplementary Figure 1G**), indicative of the faster gut transit measured in the vehicle group compared to the treated groups (**Figure 1B**). The results imply that, especially in the jejunum, the reduced gut-motility is associated with the altered microbiota composition, both of which are caused by PD medication.

Combining both the locations and treatments showed that dopamine-, pramipexole-, and 50 % of the ropinirole-treated animals cluster distant from the ileum and vehicle, while the other half of the ropinirole-treated animals cluster close to the vehicle and ileum (**Figure 3E**). Both jejunal and ileal samples from the vehicle cluster closer together (PERMANOVA for location: R^2^= 0·199, p=0·01 VH vs mean R^2^= 0·294, p=0·001 in treatment groups). Overall, the results indicate that the PD medication and dopamine-(agonist) treatments affected the small intestinal bacterial composition, which may be due to the changes in gut motility described above.

### Parkinson’s disease medication induced microbial changes mirror changes reported in patients

The ordination analyses revealed the PD treatments to have a profound effect on the microbiota composition (**Figure 3**). To identify which bacterial taxa are significantly affected by the treatment, we continued with differential abundance analysis. Focusing on the top 10 most abundant taxa in all groups, *Muribaculaceae spp*. (previously known as S24-7, most often isolated from *Murinae* species but are also found in humans ^68^) and *Lactobacillus spp*. appeared to be the most prominent members in both jejunum and ileum (**Figure 4A-B**). Using PCA (Principal Component Analysis), *Muribaculaceae* and *Lactobacillus* showed the strongest association with PC1 (explaining ~77% of the variation) and PC2 (explaining ~11·0% of the variation) respectively (**Supplementary Figure 2 A-B**). In ileum, *Lactobacillus* is associated with the same axis (PC2) that separates the treated groups from the vehicle group (**Supplementary Figure 2B**). Remarkably, in the dopamine-, pramipexole- and ropinirole-treated groups, *Lactobacillus spp*. in the ileum was significantly increased compared to the vehicle group (Dunnett’s test p=0·001, 0·003 and, 0·047, respectively) (**Figure 4B**), which is in agreement with the observation that *Lactobacillus* is associated with PC2, described above.

From the top 10 taxa in jejunum, only *Romboutsia spp*. were significantly decreased (Dunnett’s test p= 0·022) in the pramipexole-treated group compared to the vehicle (**Figure 4A**). In ileum, *Lachnospiraceae spp*. were decreased in both dopamine- and pramipexole-treated groups (Dunnett’s test p=0·033 and 0·034 respectively), while *Enterorhabdus spp*. were decreased in the dopamine-treated group, and *Allobaculum spp*. increased in the pramipexole-treated group (Dunnett’s test p=0·011, and 0·002 respectively), compared to vehicle (**Figure 4B**).

Next, we used LEfSe (Linear discriminant analysis Effect Size) ^64^ for differential abundance analysis on OTU (operational taxonomic unit) level. From the LEfSe analyses (**Figure 4C-D**), the main discriminant feature separating the vehicle from the other treatment groups were species from the *Muribaculaceae* family in both jejunum and ileum. The main discriminant feature separating the groups treated with dopamine or PD medication from the vehicle group are species from the *Lactobacillus* genus, in accordance with the results observed in **Supplementary Figure 2B** and **Figure 4B**. Species from the *Prevotellaceae, Lachnospiraceae* and *Muribaculaceae* family in jejunum or ileum were significantly decreased in almost all treated groups compared to the vehicle group. Species from the *Lactobacillus* genus were significantly increased in the ileum. Among the other significantly increased differential taxa were *Bifidobacterium* in the ileum of the pramipexole-treated group and *Enterococcus* in the jejunum of the dopamine-treated group.

Next to the LEfSe analysis the zero-inflated log-normal model (ZILM) form the metagenomeSeq R-package^60^ was used for differential abundance analysis (as recently at least two methods are recommended to gauge for concordance^69^). Although the ZILM method appeared to be more stringent, it strengthened some of the observed differential abundances observed with the LEfSe method (**Supplementary Figure 2CD, Supplementary Table 1**). Overall, these findings are highly relevant as *Prevotellaceae* and *Lachnospiraceae* taxa are often reported to be decreased, while *Bifidobacterium* and *Lactobacillus* taxa are often reported to be increased in PD patients compared to HC subjects, where the latter could interfere with the uptake of levodopa.

In particular, the increase in *Enterococcus* and *Lactobacillus* may be relevant for the levodopa decarboxylation activity in the jejunum, as species from these genera have been shown to decarboxylate levodopa and could restrict the available levels in the blood.^48^ Therefore, we tested the decarboxylation activity of levodopa (LDC) in the jejunal and ileal samples as well as the levodopa uptake in blood samples. Although, no significant differences were observed in LDC or levodopa uptake between the tested groups (**Supplementary Figure 3A-B**), there was a negative correlation (spearman correlation analyses) between only OTUs from *Lactobacillus* genus out of the top 50 most abundant OTUs (1·2 % from total) in the jejunum, and the plasma levels of levodopa/carbidopa (**Figure 4E, Supplementary Table 2**) but not the LDC (**Supplementary Figure 3C**). After FDR correction *Lactobacillus* sp. OTU_168 remained significantly correlated with the plasma levels of levodopa/carbidopa (**Figure 4E**). Collectively, these results imply an interference of *Lactobacillus* species with levodopa uptake as we previously reported.^48^

## Discussion

This study unravelled the effect of PD medication, pramipexole or ropinirole in combination with levodopa/carbidopa on the small intestinal motility and the associated alteration in the bacterial counts, in the ileum (**Figures 1, 2**). Small intestinal motility is one of the factors influencing SIBO^40^, and is prevalent in PD patients.^41–43^ PD medication has been also shown to be associated with GI symptoms^44^ and reduced transit times.^45^ Here, we show that the PD medication *per se*, in healthy rats, affect the microbial profile, besides its effect on gut motility (**Figure 3**). Dopamine, which is not used as PD treatment, as it cannot pass the blood-brain barrier, but can still be produced from levodopa endogenously by the human dopa decarboxylase (DDC) or exogenously via bacterial tyrosine decarboxylases (TDC) in the periphery ^48^, also affected the microbiota profile (**Figure 3–4**). Despite the substantial number of reports describing an effect of dopamine on gut motility ^23–36^, dopamine did not exert a significant effect on the gut motility in our study (**Figure 1**). This could be due to the metabolism during the absorption process. For example, the first-pass metabolism of dopamine is predominant in the intestine of dogs and its oral bioavailability was only approximately 3% with a half-life of 10·8 minutes.^70^ In contrast to dopamine, pramipexole and ropinirole, both of which showed a significant effect on small intestinal motility (**Figure 1**), have a much higher bioavailability with a longer half-life compared to dopamine. Pramipexole has an oral bioavailability of ~90%, a long half-life (between 11·6-14·1 hours), minimal metabolism (70-78% excreted unchanged in urine) and is primarily eliminated via the kidney.^71^ Ropinirole has an oral bioavailability of ~50%, a shorter half-life of approximately 6 hours (ranging from 2 to 10 hours) and only 10% is excreted unchanged in urine and cleared by hepatic metabolism.^72^ The observation that 8/10 of the points (up to 3^rd^ quartile) in the dopamine group are below the median of the VH group (**Figure 1**) still implies that dopamine could affect gut motility, but in a less pronounced manner compared to dopamine agonists. Importantly, gut motility contributed significantly to the variation observed in the microbiota profiles and the faster transit times (higher geometric centre) associated closely to the vehicle group (**Figure 3C-F and Supplementary Figure 1 G-H**). Moreover, both dopamine and its agonists-treated groups share similar differentially abundant taxa (**Figure 4, Supplementary Figure 2CD, and Supplementary Table 1**), implying that dopamine, pramipexole, and ropinirole act through similar mechanisms, likely by altering gut motility, and consequently, differentiating the microbiota composition.

Although the microbial profiles were altered in both jejunum and ileum, the spread of samples was larger in the jejunum compared to the ileum (**Figure 3**). This high spread in the jejunal microbiota composition upon exposure to PD medication implies that the medication has a stronger effect on the jejunal motility. This effect could be due to the rapid absorption of the drugs in the proximal small intestine^70,72,73^, resulting in the highest local drug concentration in the proximal small intestine. Plausibly, these drugs could also elicit a direct effect on the microbiota, which warrants further elucidation.

The often-reported increase in *Lactobacillus, Bifidobacterium, Akkermansia*, and the decrease in *Lachnospiraceae* and *Prevotellaceae* have been reported as a common finding among the several studies investigating the faecal gut microbiota composition between PD patients and HC subjects.^17^ Remarkably, for most of these altered taxa (except Akkermansia), a similar alteration was observed in the small intestine of the healthy rat model employed in this study (i.e., not PD model). This infers that the observed changes in microbiota composition in PD patients is, at least partly, due to the PD medication and not the disease *per se*. The alterations observed in the small intestine were in agreement with the changes in microbiota composition previously reported in the faecal samples of PD patients. This is not surprising since 85·9% of the taxa are shared between small intestine and faeces and correlate significantly (Spearman’s R=0·69, R^2^=0·48, p<2·2E-16 on log transformed data, calculated from Table S3 in ^74^) and are ultimately washed-out through the large intestine.

Intriguingly, *Lactobacillus spp*. were found to be the discriminating factor between all treated groups compared to the vehicle, in the ileum (**Figure 4D**). This is in agreement with the pooled data study of Romano *et al*.^17^, where this genus was also the most strongly enriched in PD patients. Remarkably, a significant negative correlation was observed between species from the *Lactobacillus* genus and levodopa uptake (**Supplementary Figure 2**). Several *Lactobacillus* species harbour TDC enzymes, which can decarboxylate levodopa.^48^ Thus, the observed association implies that the higher abundance of *Lactobacillus* species could have reduced the levodopa levels in the systemic circulation, potentially affecting the levodopa bioavailability. However, no positive correlations were observed between the LDC activity, which could be due to the relatively short treatment period (14 days), our activity measure was either to selective, or that other unknown factors play a role in the association with the reduced uptake observed.

Overall, this study showed the impact of commonly prescribed PD medication and dopamine on the small intestinal motility, SIBO, and microbiota composition, irrespective of the PD disease. Importantly, the microbial alterations observed in our healthy rat model in the small intestine are consistent with the faecal microbial alterations observed in human cross-sectional studies comparing PD with HC subjects. Taken together, the study highlights the urgency of taking PD medication in consideration during the assessment of the PD-associated microbiota.

## Availability of data and materials

All data generated or analysed during this study are included in this published article and its supplementary information files. The 16s rDNA metagenomic sequence data were deposited under BioProject number PRJNA725395.

## Authors’ Contributions

S.P.K. and S.E.A conceived and designed the study. S.P.K., A.B., G.D. S.E.A performed the experiments and S.P.K. and S.E.A. analysed the data. S.P.K. wrote the original manuscript that was reviewed by S.E.A and G.D. Funding was acquired by S.E.A. All authors read and approved the final manuscript.

---

**Supplementary Figure 1.**
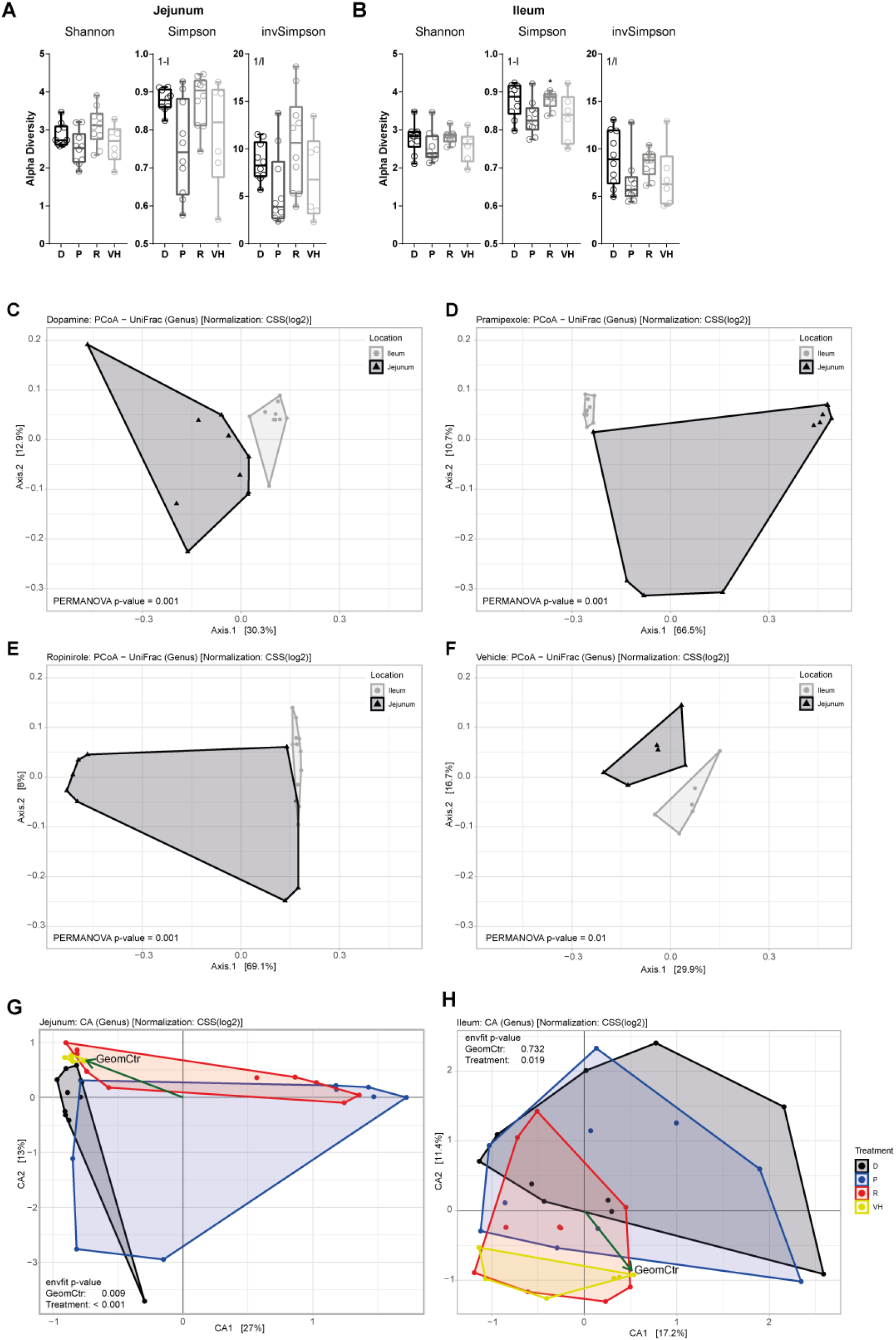
Differences between the jejunal and ileal microbiota profile within and among the treated groups. (**A,B**) represent the alpha diversity (Shannon and Simpson) of the jejunum (**A**) and ileum (**B**). Boxes represent the median with interquartile range, and whiskers represent the maxima and minima. Significance compared to VH (asterisks) was tested with One-way-ANOVA followed by Fisher’s LSD test with FDR correction. D, dopamine; P, pramipexole; R, ropinirole; VH, vehicle (10% sucrose). (**C-F**) represent a PCoA using unweighted UniFrac distances at genus level using CSS scaled data of dopamine (**C**), pramipexole (**D**), ropinirole (**E**), and vehicle (**F**) treated groups in jejunum (depicted by black triangles) and ileum (depicted by grey circles). Significant contribution of the variables to the variance of the PCoA was tested with Permutational Multivariate ANOVA (PERMANOVA). (**G,H**) depict a Correspondence Analysis (CA) at genus level using CSS scaled data of jejunum (**G**) and ileum (**H**) with arrows indicating the direction of the geometric centre towards the vehicle group. The significant contribution of the environmental vectors was tested with permutational test from *envfit* function from R package *vegan*.

**Supplementary Figure 2.**
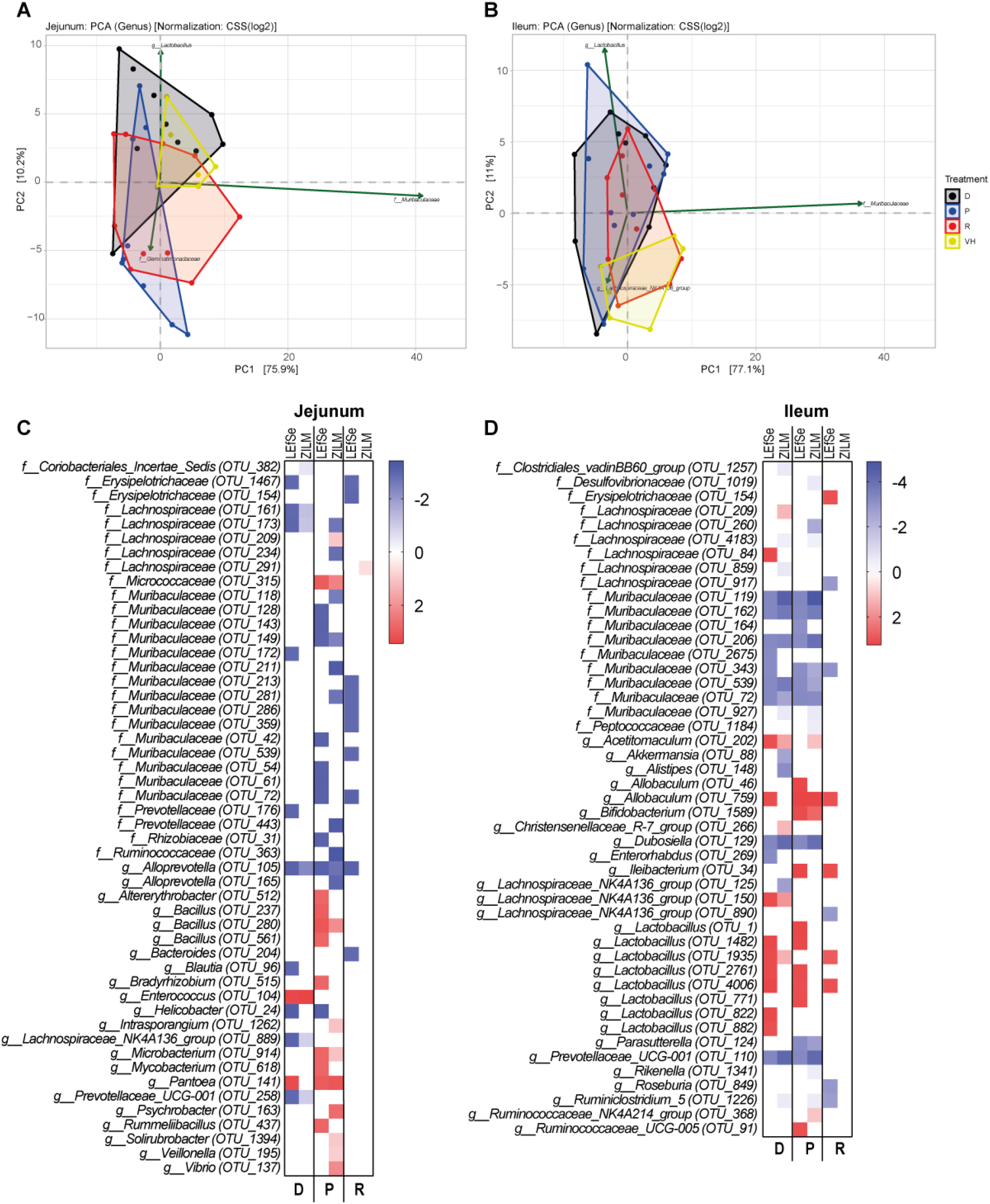
Principal component analysis and differential abundance analysis comparing LEfSe and zero-inflated log-normal model (ZILM). (**A,B**) depict a Principal Component Analysis (PCA) at genus level using CSS scaled data of jejunum (**A**) and ileum (**B**). Arrows represent the top 3 contributing genera to the variation of PC1 and PC2. (**C,D**) depicts a comparison between the LEfSe and zero-inflated log-normal model (ZILM), in jejunum (**C**) and ileum (**D**). A positive value depicts a discriminating feature to be increased from the vehicle in the LEfSe analysis and resembles the fold change for the ZILM analysis. For the LEfSe analysis significance was tested using one-way-ANOVA followed by a Kruskal-Wallis (KW) test and Linear Discriminant Analysis (LDA). A significant feature was considered when KW p-value < 0·01 and Log(LDA score) > 2. For the ZILM analysis a significant feature was considered when FDR < 0·1.

**Supplementary Figure 3.**
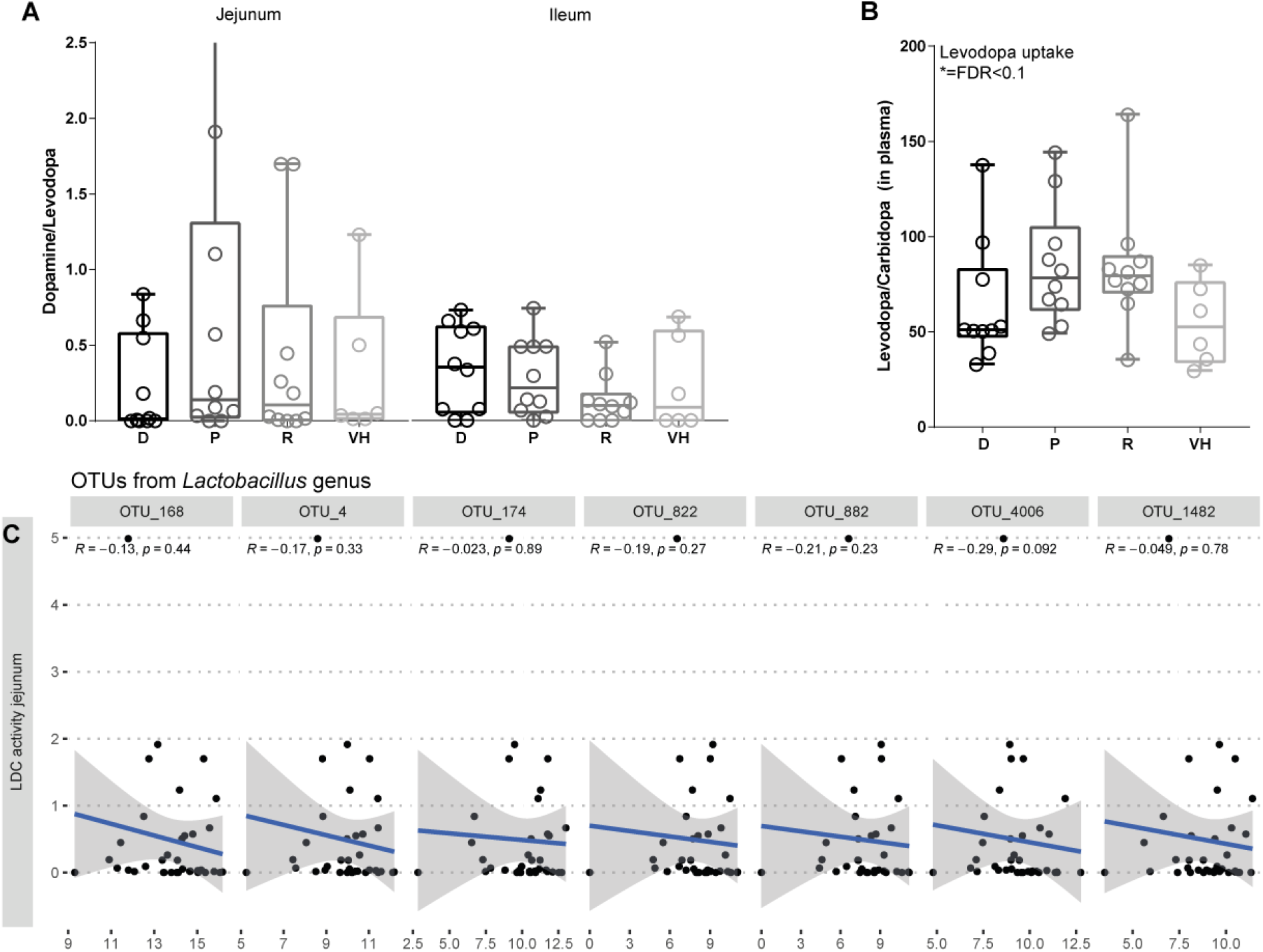
Levodopa decarboxylase activity and levodopa uptake. (**A**) depicts the levodopa decarboxylase activity (LDC) in the jejunal and ileal content. (**B**) represents the levodopa/carbidopa ratio in plasma at Cmax. D, dopamine; P, pramipexole; R; VH, vehicle (10% sucrose). Boxes represent the median with interquartile range, and whiskers represent the maxima and minima. Significance compared to VH (asterisks) was tested with One-way-ANOVA followed by Fisher’s LSD test with FDR correction. (**C**) shows graphs with linear models and spearman correlations of the significant OTUs from **Figure 4E** with the LDC activity.

